# Exosomal miR-106b-5p promotes Mtb survival via targeting CREB5 followed by SOAT1-CIDEC and CASP9-CASP3 pathway

**DOI:** 10.1101/2021.08.11.456001

**Authors:** Haotian Chen, Chonghui Li, Taohua Song, Jiao Gao, Wenjing Li, Yurong Fu, Zhengjun Yi

**Affiliations:** School of Laboratory Medicine, Key Laboratory of Clinical Laboratory Diagnostics in Universities of Shandong, Weifang Medical University, 261053 Weifang, Shandong, China; Department of Medical Microbiology, School of Basic Medicine, Weifang Medical University, 261053 Weifang, Shandong, China

**Keywords:** *Mycobacterium tuberculosis*, exosomes, miR-106b-5p, apoptosis, lipid

## Abstract

Tuberculosis(TB) is one of the top ten fatal diseases, but the research on the mechanism of TB is still not perfect. Exosome, as an important intercellular signal transmission signal vehicle and the mechanism of exosomes in the interaction between macrophages and *Mycobacterium tuberculosis*(Mtb), is crucial for TB treatment. In the study, we found that exosomes, derived from Mtb-infected macrophage, exhibited differential enrichment in different organs in mice, causing inflammatory cell infiltration in lungs. Further experiments *in vitro* showed that exosomes resulted in increased lipid synthesis and inhibition of apoptosis in normal macrophages. In order to further explore its molecular mechanism, bioinformatics analysis showed that miR-106b-5p was up-regulated in exosomes. Subsequently, we verified miR-106b-5p was increased through a large number of blood samples from TB patients. In addition, we demonstrated that miR-106b-5p was upregulated in exosomes from Mtb-infected macrophages, which can be engulfed by uninfected macrophages and further result in miR-106b-5p increase. We next found that miR-106b-5p mediated the same effect as the exosomes derived from infected macrophage. Through further research, we indicated that miR-106b-5p promoted lipid droplet accumulation through regulation of Creb5-SOAT1-CIDEC and suppressed macrophage apoptosis via Creb5-CASP9-CASP3 pathway, which ultimately led to Mtb survival. These findings provide a certain theoretical basis and ideas for the diagnosis and treatment of TB as well as the selection of biomarkers.

## Introduction

Tuberculosis(TB) is an infectious bacterial disease caused by *Mycobacterium tuberculosis*(Mtb). It spreads between humans through the respiratory route. The most common pathological changes are concentrated in lungs, but they can damage to any organization (Ravimohan et al, 2018). Vaccination and other preventive measures have a certain effect, however, TB is still the world’s leading cause of death from infectious pathogens, surpassing human immunodeficiency virus/acquired immunodeficiency syndrome (HIV/AIDS) for the first time (Gupta et al, 2018; Tornheim et al, 2018). The World Health Organization (WHO) estimates that there are approximately 10.4 million new cases and 1.8 million deaths from TB each year (Goletti et al, 2018). One third of the new cases (about 3 million) are still unknown to the health system, and many people do not receive proper treatment (Pezzella et al, 2019). Alveolar macrophages represent both the main host cell and the first line of defense against Mtb infection. Macrophage apoptosis plays a crucial role in host defense against Mtb (Lam et al, 2017). Complex lipids exist in the cell wall of Mtb, and as the main effector molecules, and actively interact with the host to regulate its metabolism and stimulate immune response. However, host cells can also determine the fate of infection by regulating lipid homeostasis (Gago et al, 2018). Macrophages can release bacteria into the extracellular environment in the form of apoptotic bodies,which can be swallowed by other macrophages and further degrading Mtb. However, Mtb can also destroy macrophage apoptosis (Abdalla et al, 2020).

Host-derived or pathogen-derived biomolecules are ideal biomarkers of TB infection. Exosomes are defined as 30-150nm-sized vesicles derived from cells that are released into body fluids and participate in intercellular communication and immune regulation. As a communication medium between cells, exosomes carry biologically active molecules including microRNA (miRNA), which can regulate gene expression and function in recipient cells (Messina et al, 2020). These vesicles have become a new platform for improving the clinical diagnosis and prognosis of different infectious diseases and cancers, and their role as alternative biomarkers for improving the diagnosis and treatment of TB has been confirmed in a large number of literatures (Hu et al, 2019; Diaz et al, 2016; Arya et al, 2020; Biadglegne et al, 2021). miRNAs are tiny non-coding ribonucleic acids, approximately 21 to 25 nucleotides in length. Researchers have proved that miRNA plays an important role in controlling gene translation or regulating gene degradation, and participates in regulating important genes during Mtb infection (Riolo et al, 2020; Schäfer et al, 2020). Functional miRNAs can be encapsulated in exosomes and delivered to recipient cells, which then cause specific regulation of their transcriptome (Alipoor et al, 2016). Studies have shown that many miRNAs are differentially expressed in exosomes released from BCG-infected macrophages (Alipoor et al, 2017). These miRNAs indicated that metabolic reprogramming may occur, which is conducive to the survival of Mtb.

Current studies have indicated that the miR-17 family plays an important role in the pathogenesis of TB, which is mainly composed of miR-17, miR-18a, miR-20a, miR-93, miR-106a, miR-106b and other miRNA. Studies have shown that miR-17-5p was increased in the serum of TB patients and is related to the survival of BCG and Mtb in macrophages (Tu et al, 2019; Kumar et al, 2016). Our previous research found that miR-18a-5p and miR-20b-5p are expressed differently in Mtb-infected macrophages, and may play a specific role in regulating the survival of Mtb (Yuan et al, 2020; Zhang et al, 2019). However, the role of miR-93 and miR-106 in TB has not been fully studied.

In this study, we explored the role of exosomes secreted by macrophages infected with Mtb. A mouse model was established to study the accumulation of exosomes in organs and tissues, as well as corresponding pathological changes of them. A macrophage co-culture model of exosomes was established to study the effects of exosomes on the survival of Mtb and macrophage phenotype. According to the corresponding phenotypic changes, miR-106b-5p was selected for subsequent studies. This study reveals the role and mechanism of miR-106b-5p in Mtb infection from the perspective of exosomes, which provides a theoretical basis for the elucidation of molecular mechanisms, follow-up related research, as well as potential biomarker and therapeutic target.

## RESULTS

### Mtb-infected macrophage-derived exosomes are differentially aggregated in organs

In order to study the function of exosomes secreted by Mtb infected macrophages, we constructed an *in vitro* model of Mtb infection of macrophages, then extracted exosomes and identified its marker protein CD63 (Figure 1A). A typical clear exosomal cup holder-like structure was observed by transmission electron microscope (Figure 1B). Nanoparticle size analysis showed that the particle size of macrophage derived exosomes decreased after infection (Figure 1C), This suggests that changes in the physical properties of exosomes may be closely related to their functions. To verify the difference in the distribution of exosomes after infection to mediate the corresponding effects, we constructed an *in vivo* mouse model of exosomes injected through tail vein. The exosome was labeled with PKH26 and imaged with a small animal living imaging system, which showed that enhanced fluorescence signals in the chest and abdomen of mice injected with exosomes secreted by infected macrophages (Figure 1D). In order to define the exosomes-enriched organs, tissue imaging after dissection showed that the lung and spleen showed consistent enhancement of the infected macrophage-derived exosomes at 12h, 36h, 72h after tail vein injection, and the most obvious was at 36h (Figure 1E). In order to detect the effect of exosomal organ enrichment, we found that inflammatory cells in the lungs of IM-Exo group were aggregated by tissue HE staining (Figure 1F). It is suggested that Mtb infection can cause macrophages to secrete exosomes with specific functions and can act on specific recipient cells.

**Figure 1.**
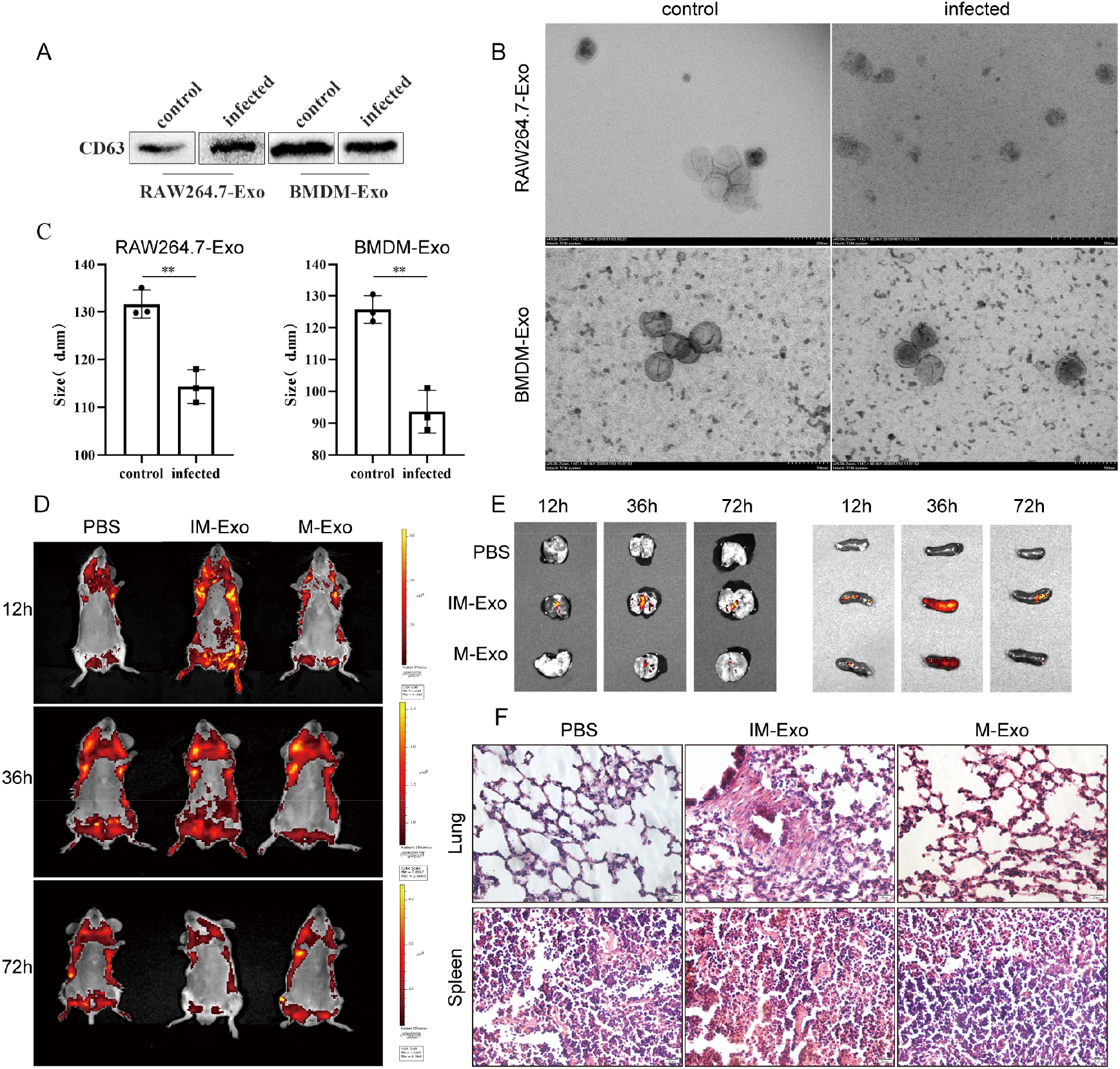
Exosomes secreted by Mtb-infected macrophages mediate specific accumulation in mouse lungs. **(A)** The corresponding exosomes were extracted from normal and infected (MOI=10) RAW264.7 and BMDM cells. Detection of the expression of exosomes marker protein CD63 by Western blot. **(B)** Ultrastructure of exosomes was observed by transmission electron microscopy. **(C)** Nanometer particle size analyzer detects the change of exosomal particle size before and after infection. **(D)** BABL /c mice were injected with PKH26 labeled exosomes through tail vein. Small animal live imaging instrument observes the difference in fluorescence on the body surface of exosomes aggregated at 12h/36h/72h. **(E)** Tissue imaging to observe the accumulation of exosomes in the lung and spleen at 12h/36h/72h. **(F)** The changes of lung and spleen cells in mice were observed by HE staining after 72 hours of tail vein injection of exosomes. \*\**p*< 0.01; IM-Exo, macrophages co-cultured with infected macrophage source exosomes; M-Exo, macrophages co-cultured with exosomes derived from normal macrophages

### Exosomes promotes Mtb survival by regulation of lipid accumulation and inhibition of apoptosis

Combined with the existing results, the regulatory effect of exosome secreted by Mtb infected macrophages on normal macrophages was studied. PKH26 was used for exosome staining and tracing to construct the macrophage model of exosome co-culture (Figure 2A). With the extension of the co-cultivation time, the fluorescence signal gradually transferred from the extracellular to the intracellular, which proved that the exosomes were gradually internalized. On this basis, the influence of exosomes on intracellular Mtb survival was discussed. According to acid-fast staining, exosomes could lead to reduced phagocytosis of macrophages, but there was no significant difference between the IM-Exo group and the M-Exo group (Figure 2B). It was observed that the colony count of IM-Exo co-cultured macrophages increased after inoculation (Figure 2C). These observations indicate that exosomes secreted by infected macrophages can promote the survival of Mtb in macrophages after infection.

**Figure 2.**
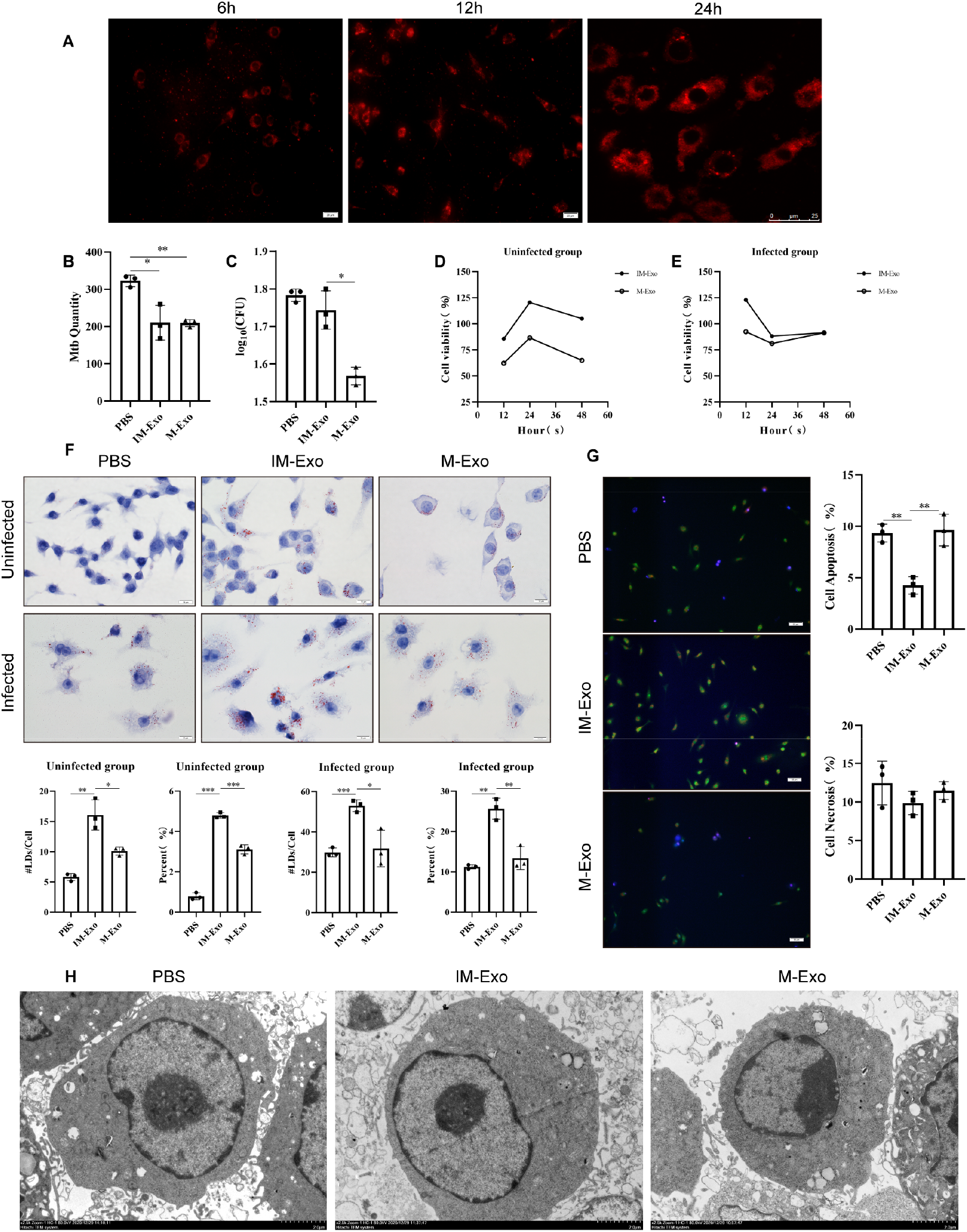
Exosomes secreted by Mtb-infected macrophages may promote Mtb intracellular survival by affecting normal cell lipid metabolism and inhibiting apoptosis. **(A)** The PKH26-labeled exosomes filtered by dye solution were co-cultured with macrophages, and the 6h/12h/24h co-cultured cells were imaged by fluorescence microscope and confocal microscope. **(B)** PBS solution, IM-Exo, and M-Exo were used to co-culture macrophages, respectively. 24h later, Mtb (MOI=10) was used to infect macrophages. Detect the amount of Mtb by acid-fast staining. **(C)** The survival rate of Mtb was measured by colony count. **(D**,**E)** CCK-8 was used to detect the viability of co-cultured macrophages and co-cultured reinfected macrophages. **(F)** Using Oil Red O to detect the changes of Lipid Metabolism in macrophages. **(G)** After whole cells were stained with AO-EB, apoptotic signals were detected by Hoechst/PI staining. **(H)** Using electron microscope to detect macrophage apoptosis.

We further investigated the effect of exosomes on macrophages. Among the co-cultured normal macrophages, the vitality of macrophages in IM-Exo group was enhanced compared with M-Exo group (Figure 2D). However, there was no significant difference in the viability of the macrophages after the co-cultured macrophages were infected with Mtb,(Figure 2E). Oil red O staining showed that in the co-cultured normal macrophages, compared with the PBS group and the M-Exo group, the average lipid droplet content of the IM-Exo group increased, and the percentage of the lipid droplet to the cell area increased. Similar results were obtained when co-cultured macrophages were infected with Mtb (Figure 2F). This suggests that exosomes released by infected macrophages promote the accumulation of lipid droplets in the recipient macrophages. Hoechst/PI staining showed that IM-Exo could inhibit apoptosis in BMDM cells, but had no significant effect on cell necrosis (Figure 2G). Agglutination and marginalization of nuclear chromatin were observed in the M-Exo group, while the PBS and IM-Exo groups were relatively normal (Figure 2H). This indicates that exosomes secreted by infected macrophages may influence lipid metabolism and inhibit apoptosis of peripheral normal macrophages by paracrine, thus mediating intracellular viability of Mtb.

### miR-106b-5p was increased in exosomes and mediated its direction change in recipient macrophages

It has been identified that exosomes contain large amounts of mRNA and miRNA, and regulate receptor cells after they can be absorbed by adjacent or distal cells. The difference between exosomes miRNA patterns and parental cells suggests that miRNA may be the key molecule for exosome to play a major role. In order to further explore the deep mechanism of the role of exosomes, we analyzed the clinical serum exosomes database by bioinformatics, and screened the differentially expressed miRNA by volcano map and heat map, in which miR-106b-5p and other genes showed stable and high expression (Figures 3A and 3B). The miRNA-mRNA interaction network is constructed and key nodes in the network diagram are analyzed (Figures 3C and 3D). It was found that miR-106b-5p have the highest number of nodes, and this suggested that it might plays a major role in the exosomes of patients with Mtb infection. Through GO (gene ontology) and KEGG (Kyoto Encyclopedia of Genes and Genomes) enrichment analysis, miR-106b-5p is involved in PI3K-Akt, MAPK, TB and other key pathways (Figures 3E-3G). By analyzing the PPI (Protein-Protein Interactions) network, we obtained the key genes, including CREB5 (cAMP-response element binding protein 5), that may be involved in the regulation of miR-106b-5p(Figure 3H).

**Figure 3.**
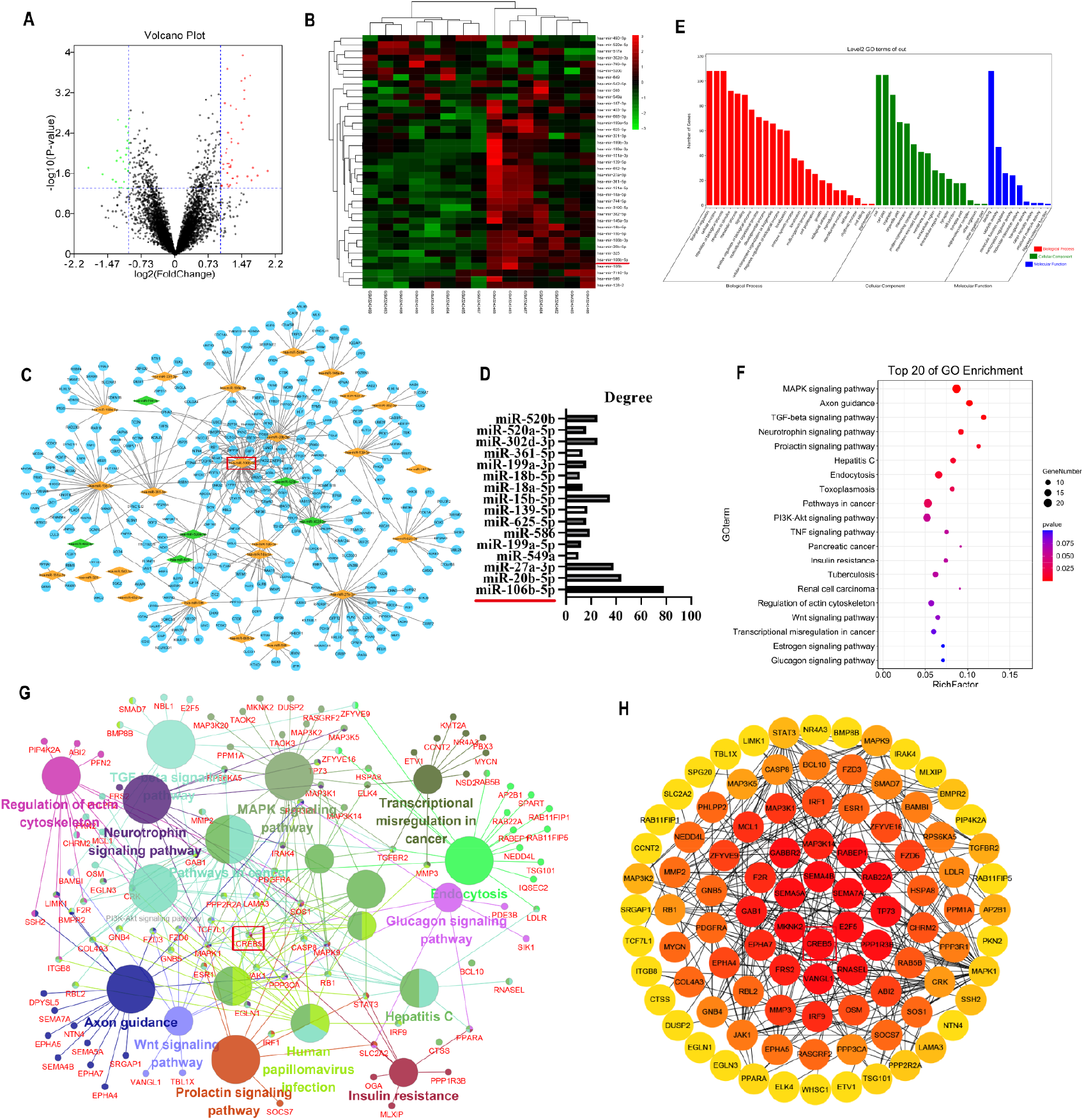
Bioinformatics analysis of miRNA specifically expressed in exosomes. **(A)** Screening differentially expressed miRNA with *p* < 0.05 and | logFc | > 1 as the standard. **(B)** Perform heat map analysis on differential miRNAs. **(C)** Cytoscape was used to map the interaction networks of differential miRNAs and their target genes. **(D)** Analyse key nodes in the interaction network using Cytoscape’s Centiscape plug-in. **(E**,**F)** Use DAVID to perform GO(**e**) and KEGG(**f**) analysis of differentially expressed genes. **(G)** Using Cytoscape to build visual KEGG network. **(H)** The PPI network was built using String and visualized with Cytoscape.

A large number of samples were used to verify the accuracy of bioinformatics analysis. Increased expression of miR-106b-5p was found in exosomes from serum and alveolar lavage fluid of PTB (pulmonary tuberculosis) patients, as well as Mtb infected macrophages *in vitro* (Figures 4A-4C). Further study found that miR-106b-5p in serum of patients with PTB also increased to some extent (Figure 4D). The ROC curve suggests that it has possibility to be a biomarker of TB (Figure 4E). At the same time, we found that the expression of miR-106b-5p in macrophages infected with Mtb was dose-dependent and time-dependent (Figures 4F and 4G). These results suggest that miR-106b-5p may plays an important role in Mtb infection. In follow-up experiments, we confirmed that infected macrophage-derived exosomes can increase the expression of miR-106b-5p in receptor cells after co-culture of normal macrophages (Figure 4H). This suggests that parent cells may regulate the expression of miR-106b-5p in receptor cells through exosomes, thus affecting the function of receptor cells.

**Figure 4.**
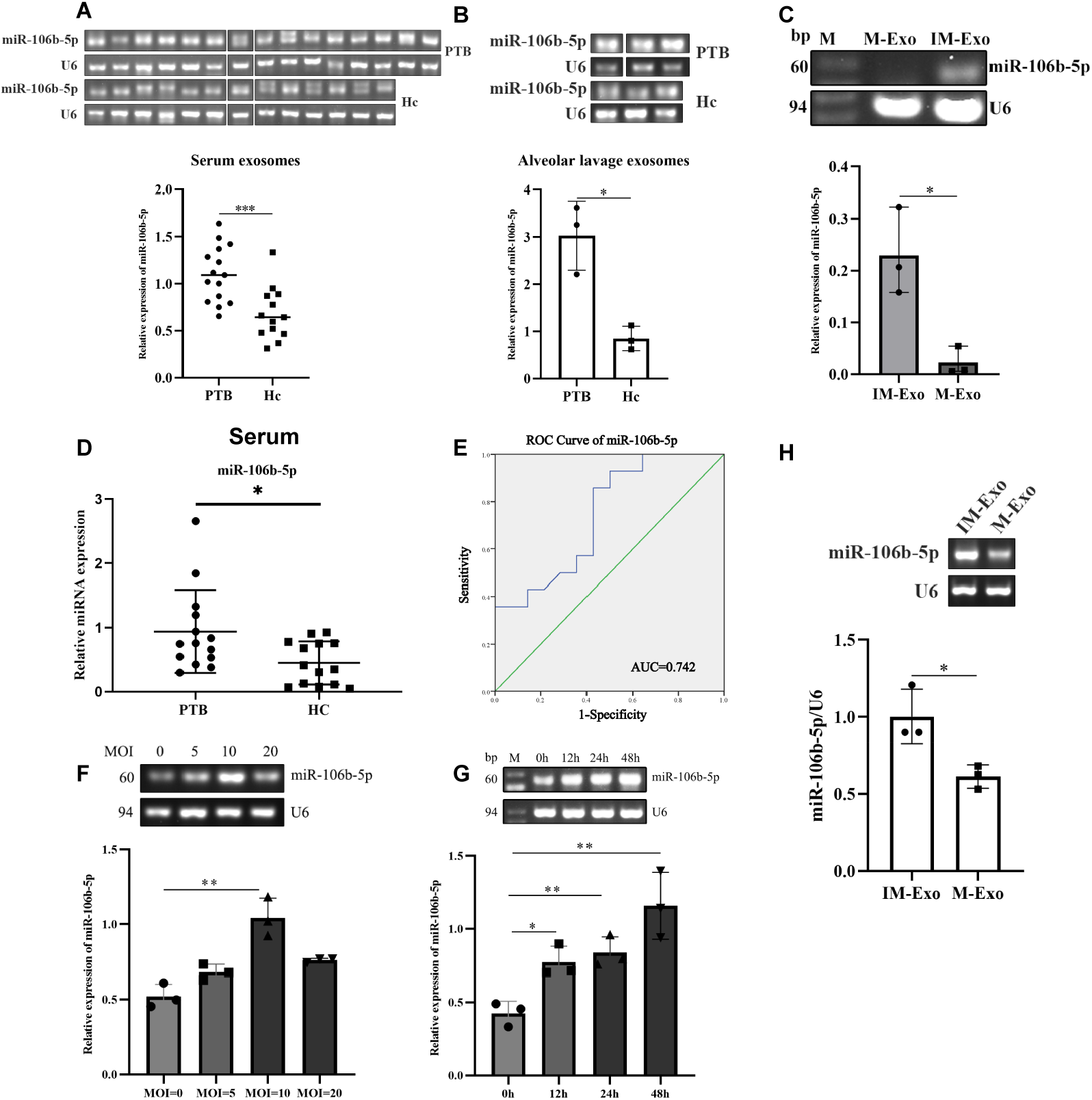
Verification of miR-106b-5p expression in exosomes by RT-PCR. **(A**,**B)** Serum exosomes and bronchoalveolar lavage fluid exosomes from patients with PTB were extracted and miR-106b-5p expression was detected by RT-PCR. **(C)** The macrophage model of Mtb infection (MOI=10) was established. The supernatant exosomes was extracted and the expression of miR-106b-5p was detected by RT-PCR. **(D)** Serum RNA was extracted from PTB patients and the expression level of miR-106b-5p was detected by qRT-PCR. **(E)** The ROC curve of miR-106b-5p was drawn. **(F**,**G)** After the bacterial quantity and time gradient models of Mtb-infected macrophages were constructed, RNA of the corresponding cells was extracted, and the expression of miR-106b-5pwas detected by RT-PCR. **(H)** The model of exocrine co-cultured macrophages was constructed. RNA, was extracted and miR-106b-5p expression was detected by RT-PCR.

### Increased miR-106b-5p induced Mtb survival by inhibiting apoptosis and promoting lipid accumulation

We further discussed the effect of miR-106b-5p regulation on intracellular Mtb survival. The *in vitro* model of macrophages with overexpression and inhibition of miR-106b-5p expression was constructed by mimics and inhibitor, and fluorescence showed that transfection efficiency was more than 80%. RT-PCR results showed that miR-106b-5p mimics significantly up-regulated the expression of miR-106b-5p. The transfection efficiency detection Figure is not shown. Subsequently, we used acid-fast staining and colony counting to verify whether miR-106b-5p had an effect on intracellular Mtb survival. Acid-fast staining showed that there is no significant change in the number of swallowing bacteria in macrophages after the regulation of miR-106b-5p expression (Figure 5A). Colony counts suggest that up-regulating the expression of miR-106b-5p promotes the intracellular survival of Mtb (Figure 5B).

**Figure 5.**
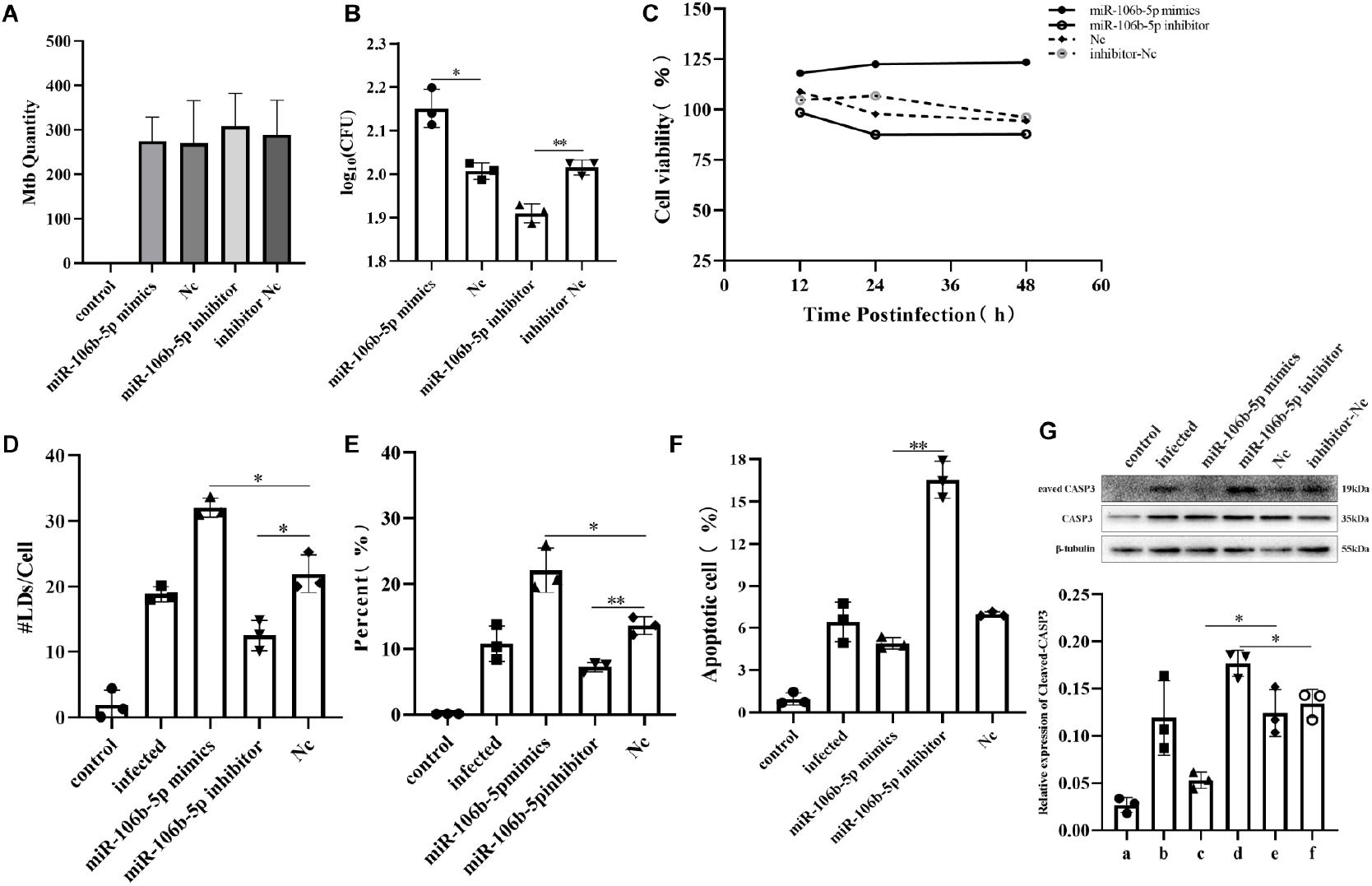
The effects of miR-106b-5p on Mtb and macrophages. **(A)** After the regulation of miR-106b-5p expression model was constructed, macrophages were infected with MOI=10. Acid-fast staining was used to count the number of swallowing bacteria in macrophages. **(B)** Intra-macrophage Mtb survival was counted using colony count(**b**). **(C)** CCK-8 was used to detect the activity of macrophages after the regulation of miR-106b-5p. **(D**,**E)** Oil red O staining was used to detect the average lipid droplet content of cells and the percentage of lipid droplets to the cell area. **(F)** Fluorescence staining was used to detect the percentage of apoptosis of macrophages. **(G)** Expression level of Cleaved CASP3 was detected by Western blot.

Then we studied the effect of miR-106b-5p on Mtb infected macrophages. The overexpression of miR-106b-5p can enhance the activity of macrophages (Figure 5C). In terms of lipid metabolism, up-regulation of the miR-106b-5p expression promoted lipid accumulation, leading to an increase in the average lipid droplet content of macrophages and percentage of lipid droplets in the cell area (Figures 5D and 5E). Then we found that apoptosis was inhibited after miR-106b-5p up-regulated, which was beneficial for intracellular survival of Mtb (Figures 5F and 5G). Combined with the phenomenon that exosomes can regulate the expression of miR-106b-5p in macrophages, the above experimental results suggest that miR-106b-5p is the key molecule for the role of Mtb-infected macrophage exosomes.

### miR-106b-5p regulates SOAT1-CIDEC and CASP9-CASP3 through targeting CREB5

We attempted to further explore the potential mechanism of miR-106b-5p promoting bacterial growth. Confirmed by research and literature, combined with the results of bioinformatics analysis, we found that Creb5, as a potential target of miR-106b-5p predicted by the four databases of TargetScan, miRWalk, miRanda and miRDB, is not only involved in the regulation of apoptosis pathway, but also related to lipid metabolism. Therefore, we verified the interaction relationship between miR-106b-5p negatively regulating Creb5 by RT-PCR (Figure 6A). Subsequently, we constructed a cell model that interfered with Creb5 expression (Figure 6B), and then verified the lipid metabolism promotion (Figures 6C and 6D) and apoptosis inhibitory effects of interference with its expression in the model (Figure 6E).

**Figure 6.**
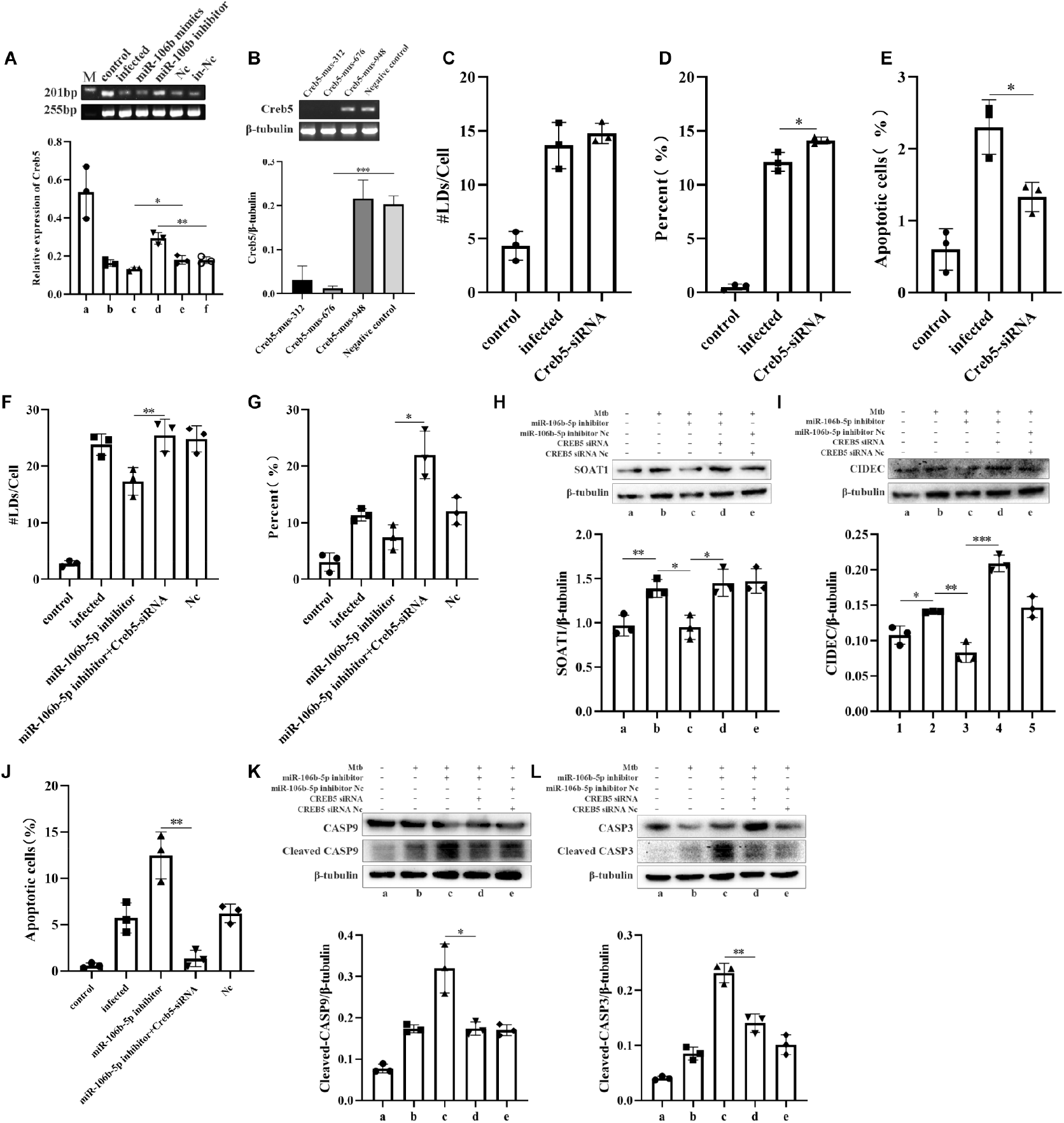
miR-106b-5p depends on Creb5 to regulate the changes of lipid metabolism and apoptosis. **(A)** n *in vitro* model was established to regulate the expression of miR-106b-5p, and the expression of Creb5 was verified by PT-PCR. **(B)** An *in vitro* model was established to interfere with Creb5 expression, and the effect of interference was evaluated by RT-PCR. **(C**,**D)** Constructing a Creb5 interference expression model, and detecting the average lipid droplet content of cells and the percentage of lipid droplets to the cell area by oil red O staining. **(E)** The apoptotic percentage of macrophages was detected by fluorescence staining. **(F**,**G)**The interference expression models of miR-106b-5p and Creb5 were constructed. Oil red O staining was used to detect the average lipid droplet content of cells and the percentage of lipid droplets to the cell area. **(H**,**I)** The expression of SOAT1 and CIDEC was detected by Western blot. **(J)** The apoptotic percentage of macrophages was detected by fluorescence staining. **(K**,**L)** The expression levels of Cleaved caspase-9 and Cleaved caspase-3 were detected by Western blot.

In order to determine whether the effect of miR-106b-5p depends on regulation of Creb5, a dual interference model of miR-106b-5p and Creb5 was constructed. In oil red O staining, the interference expression of Creb5 blocked the lipid suppressive effect of miR-106b-5p inhibitor (Figures 6F and 6G). Suppressant effect of this lipid is manifested in the negative regulation of Creb5 on SOAT1, which leads to the decrease of CIDEC and affects lipid accumulation (Figures 6H and6I). In Hoechst/PI staining, the interference of Creb5 blocked the apoptosis promoting effect of miR-106b-5p inhibitor (Figure 6J). The effect of promoting apoptosis is that Creb5 positively regulates the activation of caspase-9, which leads to the enhancement of caspase-3 activation and leads to apoptosis (Figures 6K and 6L). These results suggest that exosomes-derived miR-106b-5p positively regulates the intracellular survival of Mtb by regulating the downstream target gene Creb5 to mediate lipid accumulation and apoptosis.

## DISCUSSION

Because exosomes are relatively conservative in structure and play an important role in transmitting information from primitive cells to recipient cells, more and more studies have focused on the application of exosomes as therapeutic agents, drug delivery vehicles, and diagnostic targets (Lorenc et al, 2020). Studies have confirmed that the specific metastasis and enrichment of exosomes have an important impact on cancer progression (Mo et al, 2021). However, there are no relevant reports on the differences in the distribution of exosomes caused by intracellular bacterias infections such as TB. After testing, we found that the infected macrophage-derived exosomes have a relatively smaller particle size. We further constructed a live imaging model of exosomes and found that Mtb-infected macrophage-derived exosomes mediated higher pulmonary aggregation and led to a certain pulmonary inflammatory response, suggesting that they may play an important role in TB infection and the pathogenesis of TB.

In a variety of diseases, macrophages secrete specific exosomes to transmit specific genes, proteins or lipids and affect the immune response of related cells, thereby mediating the body’s immunity and disease progression (Lan et al, 2019). The infection of viruses, parasites, fungi and bacteria will affect the composition and function of exosomes produced by host cells, and play a role in promoting or suppressing host immunity (Schorey et al, 2015). The infected macrophage-derived exosomes were co-cultured with the recipient cells, and colony count showed that they promoted the survival of intracellular Mtb. And the foaming of macrophages and the obstruction of apoptosis are all cell changes related to Mtb survival. On this basis, it was found that exosomes derived from co-culture of macrophage on recipient cells after infection did lead to increased activity of recipient cells, vigorous lipid metabolism and decreased apoptosis.

Many mRNAs and miRNAs have been identified in exosomes, which can be taken up by adjacent or distant cells and subsequently regulate recipient cells (Sun et al, 2018). After bioinformatics analysis and verification with a large number of clinical samples, we confirmed that miR-106b-5p has high expression levels in exosomes from Mtb-infected macrophages. The miR-17 family plays an important role in TB. As a member of this miRNA family, miR-106b-5p in TB has important research value. In addition, Creb5, which appears many times in enrichment, is a target gene of miRNA predicted by 4 databases. It occupies a core position in the KEGG pathway enrichment and PPI network. And related literature points out Creb5’s potential relationship with apoptosis and lipid metabolism, suggesting that it may play a role as a downstream target gene of miR-106b-5p (Ghorpade et al, 2012; Vilà-Brau et al, 2013). In this study, we verified the targeted interaction between miR-106b-5p and Creb5 and their effects on macrophage apoptosis and lipid metabolism at the level of cell morphology and protein mechanism.

Host lipid droplets (LDS) are critical to several intracellular bacterial pathogens and viruses, and fatty acids (FAs) released from host LDS are used by the pathogens as the basis for energy and membrane synthesis (Barisch et al, 2017). Studies have found that Creb5 positively regulates the expression of SOAT1 and CIDEC by down-regulating its own expression, and may positively regulate lipid metabolism through the SOAT1/CIDEC pathway to complete lipid droplet accumulation. In addition, we also found that miR-106b-5p down-regulated the activation of CASP9, a key molecule of apoptosis, through Creb5, and then decreased the activation of CASP3, thereby negatively regulating the occurrence of apoptosis, which was similar to the effect of Creb family on apoptosis in BCG-infected macrophages (Ghorpade et al, 2012). However, the complex interaction between lipid metabolism and apoptosis still needs to be further explored.

In summary, during Mtb infection, physical and chemical properties and contents of exosomes secreted by macrophages are changed, which leads to changes in tissue enrichment. Paracrine leads to differential expression of miR-106b-5p in macrophages. Highly expressed miR-106b-5p targets the inhibition of Creb5 expression, thereby negatively regulating the expression of CASP9 and CASP3, and positively regulating the expression of SOAT1 and CIDEC. These further lead to weakened macrophage apoptosis and increased lipid droplet accumulation, which mediates the intracellular survival of Mtb. This study provides a certain theoretical basis and ideas for the intracellular survival mechanism of Mtb, the diagnosis and treatment of TB, and the selection of biomarkers.

## MATERIALS AND METHODS

### Specimen collection and ethics

Clinical samples were collected from Weifang Respiratory Disease Hospital, and relevant research was approved by the ethics committee of Weifang Medical Universtiy.

### Cell

RAW264.7 macrophages were purchased from the Cell Bank of the Chinese Academy of Sciences, and cultured in RPMI 1640 medium (Gibco) supplemented with 10% fetal bovine serum in an incubator containing 5% CO_2_ at 37°C. SPF-grade BALB/c healthy mice were purchased from Jinan Pengyue Experimental Animal Breeding Co, Ltd, 6-8 weeks old mouse tibia cells were extracted and cultured in RPMI 1640 containing macrophage colony stimulating factor (M-CSF, peprotech). The base was cultured for 7 days at a constant temperature of 37°C in an incubator containing 5% CO_2_.

### Mtb infects macrophages

We use a 1 ml syringe to pipette the Mtb strain H37Rv grown on Lowenstein-Jensen (LJ) medium into a single bacterial suspension, which was further verified by acid-fast staining. Macrophages were inoculated into a cell culture plate according to an appropriate number and cultured overnight at 37°C in an incubator containing 5% CO_2_, and cultured for a corresponding period of time at a certain multiplicity of infection.

### Preparation of exosome-free medium

Fetal bovine serum is naturally thawed at 4°C, and the complement is inactivated in a water bath at 56°C for 30 minutes. The inactivated serum is added to a sterile rigid ultracentrifuge tube. After strictly balancing the sample, centrifuge it for 16h with ultracentrifuge (CP80-WX, HITACHI, Japan) 120000g. Exosome-free culture medium was prepared from serum prepared by ultracentrifugation, and filtered by 0.22μm microporous membrane (Millipore, Billicera, MA, USA).

### Exosome isolation

The cell supernatant is collected into a centrifuge tube, 3000g, 4°C, centrifuge for 15min, take the supernatant filter with a 0.22μm microporous membrane. After all is collected, we follow the exoQuick-Tc (System Biosciences, USA) standard operating procedure to extract exosomes from the cell supernatant. Exosomes of serum samples were enriched using exoQuick (System Biosciences, USA). The extracted exosomes were detected by transmission electron microscopy, particle size analyzer and Western blot.

### Exosomes tracing

Under dark conditions, PKH26 (MaoKang Biotechnology, China) is add to Diluent C and mix gently. We then mix exosomal suspension with the same amount of Diluent C mixture gently, and let them stand for 5 minutes and pipette carefully every 1 minute, finally add an equal volume of 1% BSA to stop staining.

### *In vitro* tracing of exosomes

BABL/c mice aged 6-8 weeks were injected with PKH26 traced exosomes through the tail vein to construct an *in vitro* model. After mouse chest and abdomen hair were shaved, mouse were anesthetized according to body weight, and then living mouse and mouse organs were imaged and detected using the small animal *in vivo* imaging system (PerkinElmer, USA). The mouse organs were fixed with formalin, followed by dehydration, embedding, paraffin section and HE staining observation.

### siRNA transfection

One day before transfection, macrophages were seeded in a 24-well plate at 10×10^4^/well, and each well was transfected with 1.5 μL siRNA (20 nM) and 2 μL Lipofectamine 2000 (Thermo Fisher Scientific, USA), and Nc-siRNA was used as a transfection control. After transfection for 24h, Mtb (MOI=10) was used for infection, After infection for 24h, cells were collected for further analysis. siRNA oligonucleotides were synthesized by Jima Pharmaceutical Technology Co, Ltd. (Shanghai, China) with the sequences as follows:

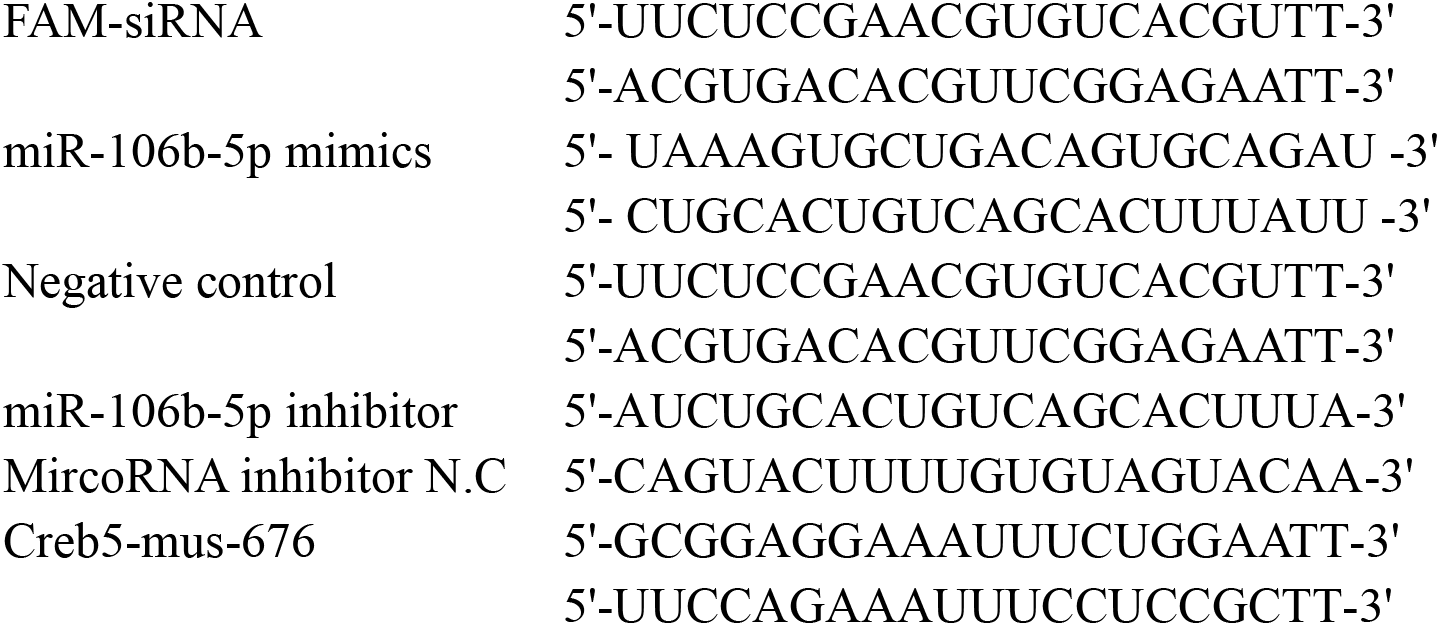

### RNA isolation and RT-PCR analysis

Trizol (invitrogen, USA) was used to extract total RNA from macrophages and exosomes follow the manufacturer’s instructions. The first-strand cDNA synthesis kit (TAKARA, Japan) and Takara Taq enzyme were used to perform RT-PCR reaction to amplify the gene sequence. The primer sequence and amplification conditions can be provided on request.

### Western blotting analysis

The cells were lysed using a cell lysis buffer (SolarBio, China) according to the instructions, and the protein concentration was determined using the BCA protein quantification kit (Beyotime, China). The protein samples were electrophoresed in sodium dodecyl sulfate-polyacrylamide gel electrophoresis (SDS-PAGE), and transferred to PVDF membrane, blocked with 5% skim milk. After incubation with corresponding primary antibody, it was incubated with HRP-labeled goat anti-rabbit IgG secondary antibody.

### Antibodies

Polyclonal rabbit anti-caspase-9(9504S), anti-caspase-3(9662S)were acquired from Cell Signaling Technology (Beverly, MA, USA). Anti-CIDEC(PA1-4316)was purchased from Thermo Fisher Scientific. Polyclonal antibody anti-SOAT1 was purchased from Abcam (Cambridge, UK). Rabbit anti-CD63 antibody(D260973), Rabbit -anti-β-tubulin(D110022)and HRP-labeled goat anti-rabbit IgG secondary antibody(D110058)were purchased from(Shanghai, China).

### Statistical analyses

Statistical analysis software IBM SPSS Statistics 22.0 was used for statistical analysis, t-test was used for pairwise comparison, single-factor analysis of variance was used for multiple comparisons between groups, *P*<0.05 was considered statistically significant.

## DATA AVAILABILITY

The data sets used for the current study are available from the corresponding author upon reasonable request.

## ACKNOWLEDGEMENTS

This work was supported by the Major Program of Shandong Province Natural Science Foundation of China (No. ZR2018ZC1054). The funder had no role in the design of the study and collection, analysis, and interpretation of data and in writing the manuscript. We thank Ziyang Liu, Xiangjuan Zhang, Nana Song, Kunshan Gao, and Heng Li for their technical support.

## AUTHOR CONTRIBUTIONS

HT.C, YR.F. and ZJ.Y. conceived and supervised the study. YR.F, HT.C. participated in the study design, analyzed the data, and wrote the manuscript. HT.C, CH.L, J.G, TH.S, WJ.L. performed the experiments. HT.C. and YR.F. provided helpful discussion and comment on study results. All authors read and approved the manuscript.

## Supplementary Information

**Figure 5-figure supplement 1.**
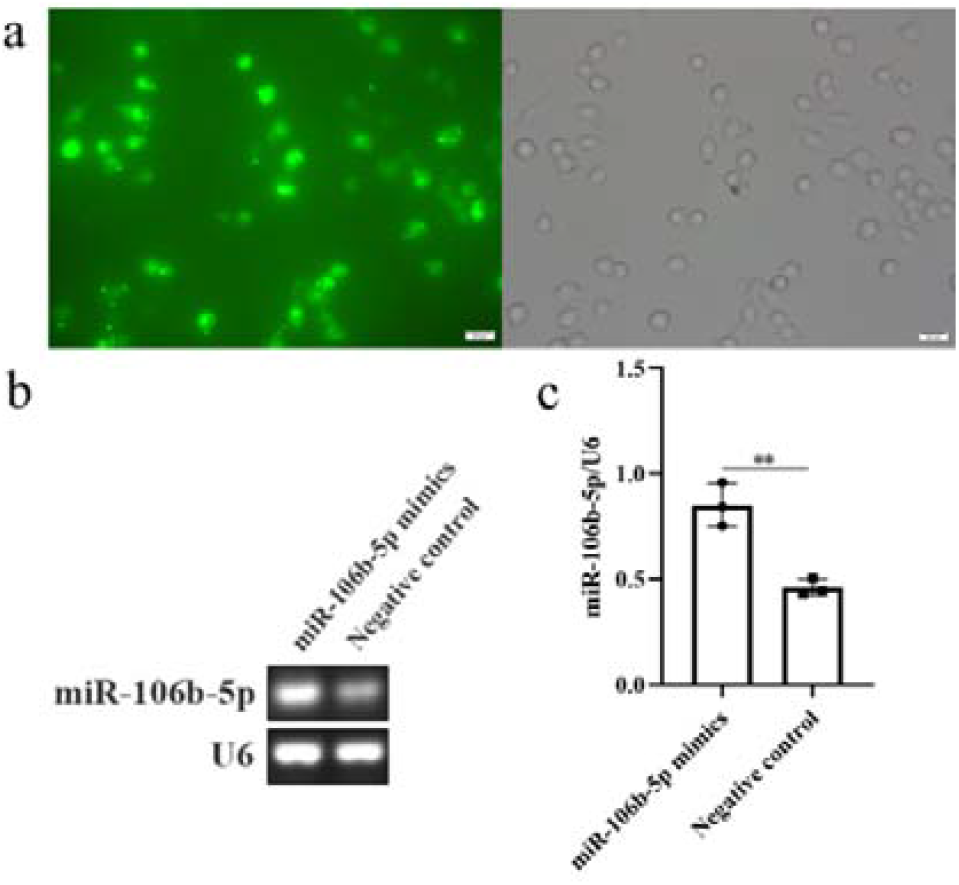
Verification of transfection effect. **a** The transfection efficiency was more than 80% under fluorescence microscope. **b**,**c** The transfection efficiency of miR-106b-5p mimics was verified by RT-PCR.

**Figure 5D, 5E-figure supplement 1.**
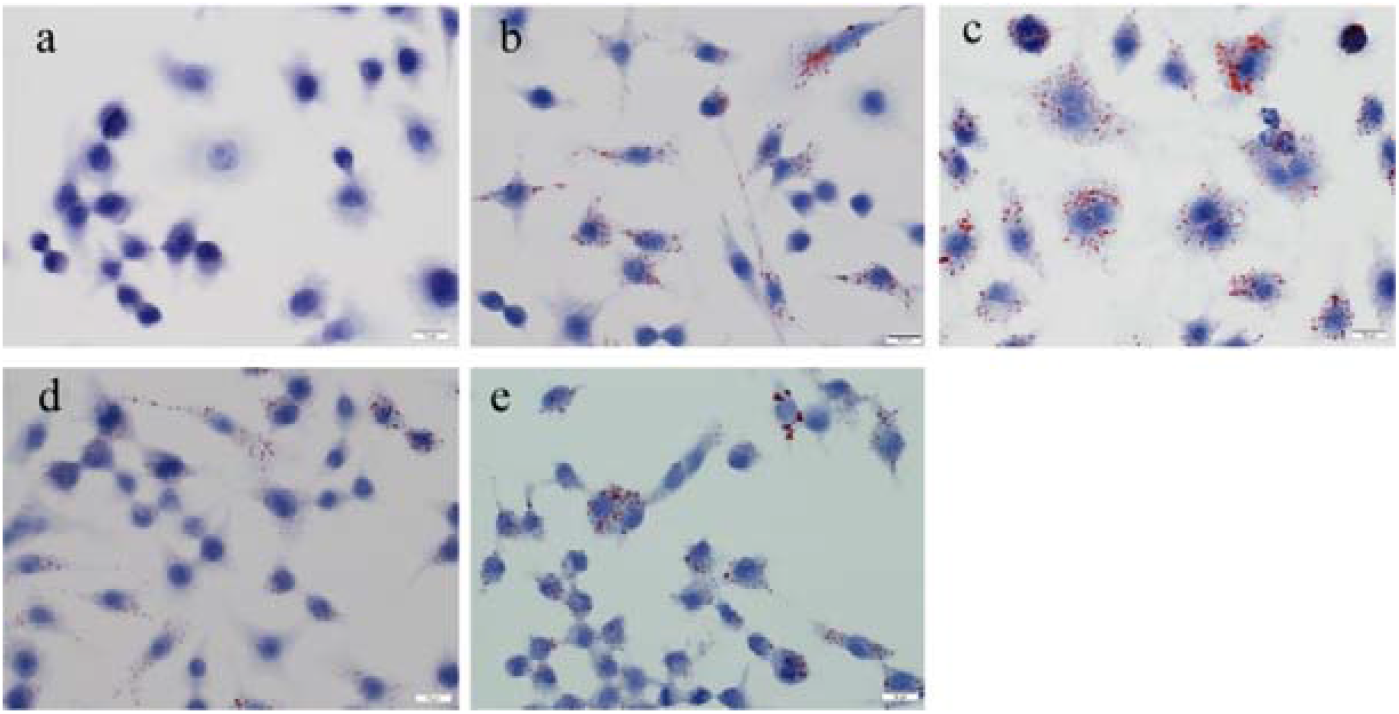
Detection of lipid changes in oil red O after regulation of miR-106b-5p **a** normal group; **b**: infection group; **c**: miR-106b-5p mimics group; **d**: miR-106b-5p inhibitor group; **e**: Nc group

**Figure 5F-figure supplement 1.**
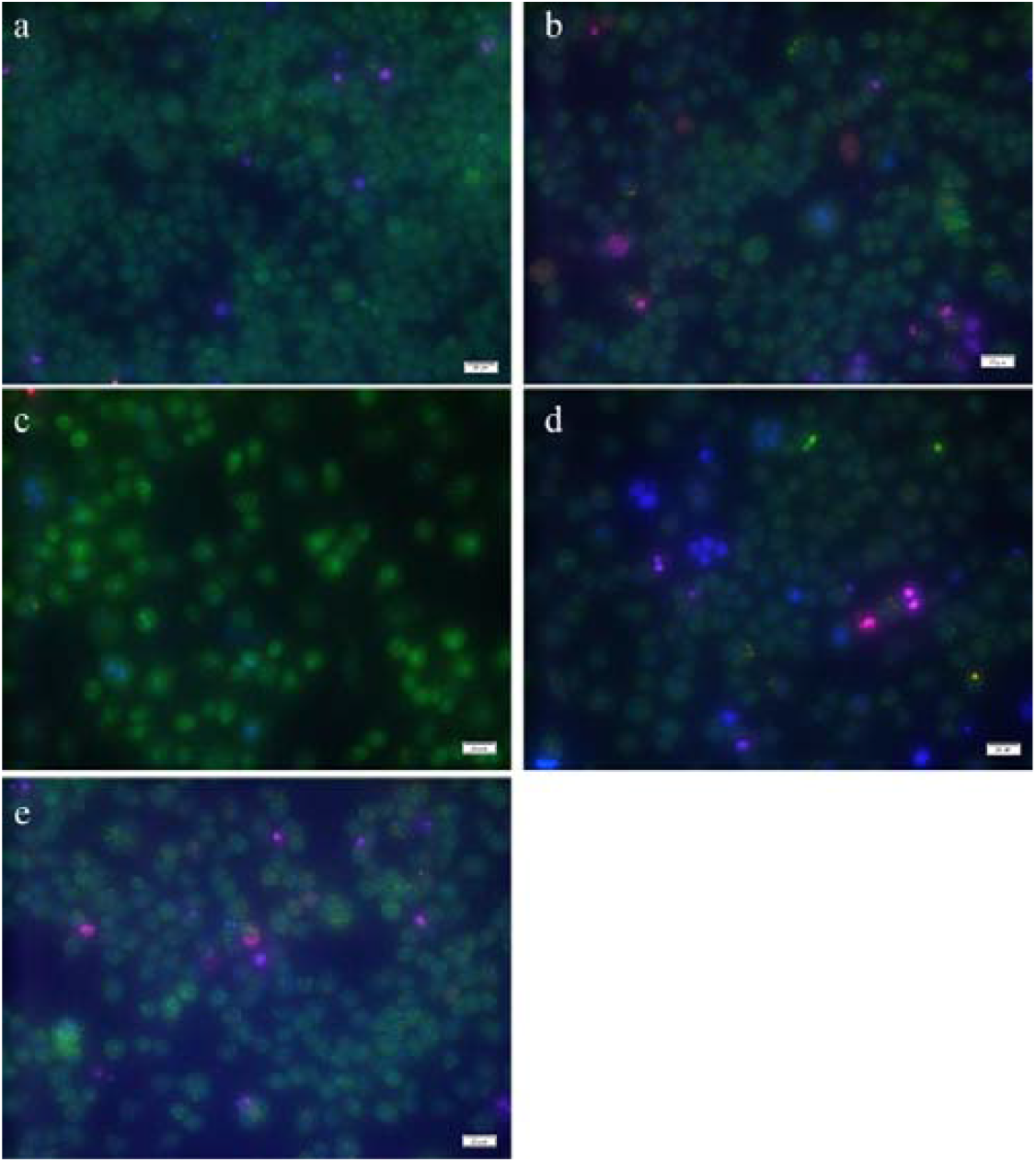
Effects of miR-106b-5p regulation on cell apoptosis **a**: normal group; **b**: infection group; **c**: miR-106b-5p mimics group; **d**: miR-106b-5p inhibitor group; **e**:Nc group

**Figure 6C, 6D-figure supplement 1.**
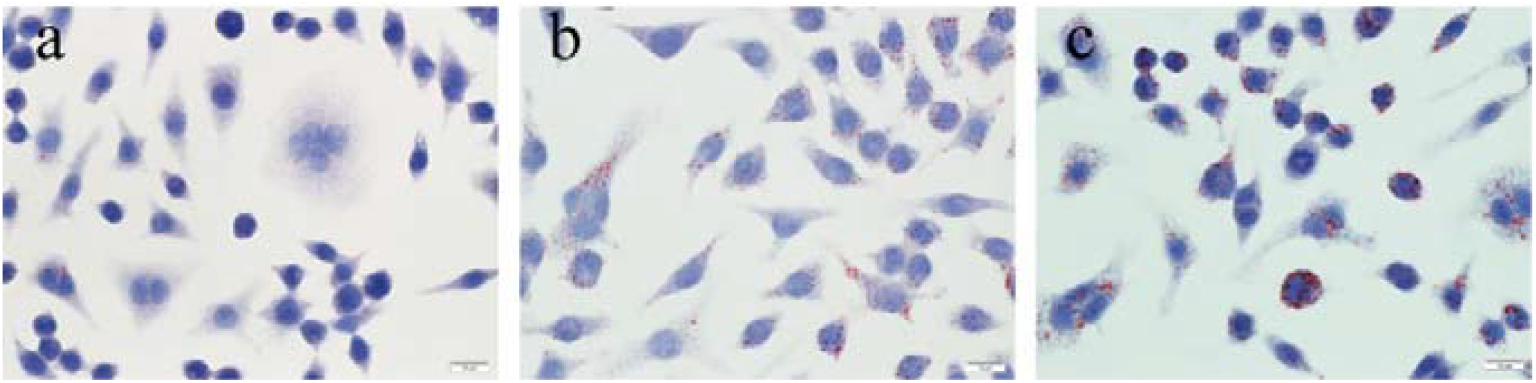
Lipid changes detected by oil red O after Creb5 regulation **a**: Normal group; **b**: Infection group; **c**: Creb5-siRNA group

**Figure 6E-figure supplement 1.**
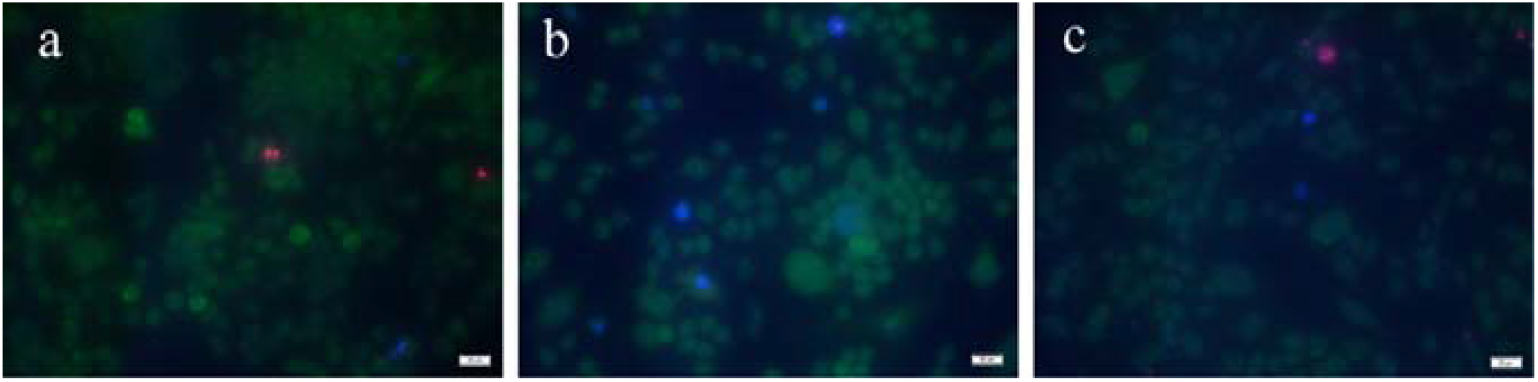
Detection of apoptosis by fluorescence staining after regulation of Creb5 **a**: normal group; **b**: Infection group; **c**: CREB5-siRNA group

**Figure 6F, 6G-figure supplement 1.**
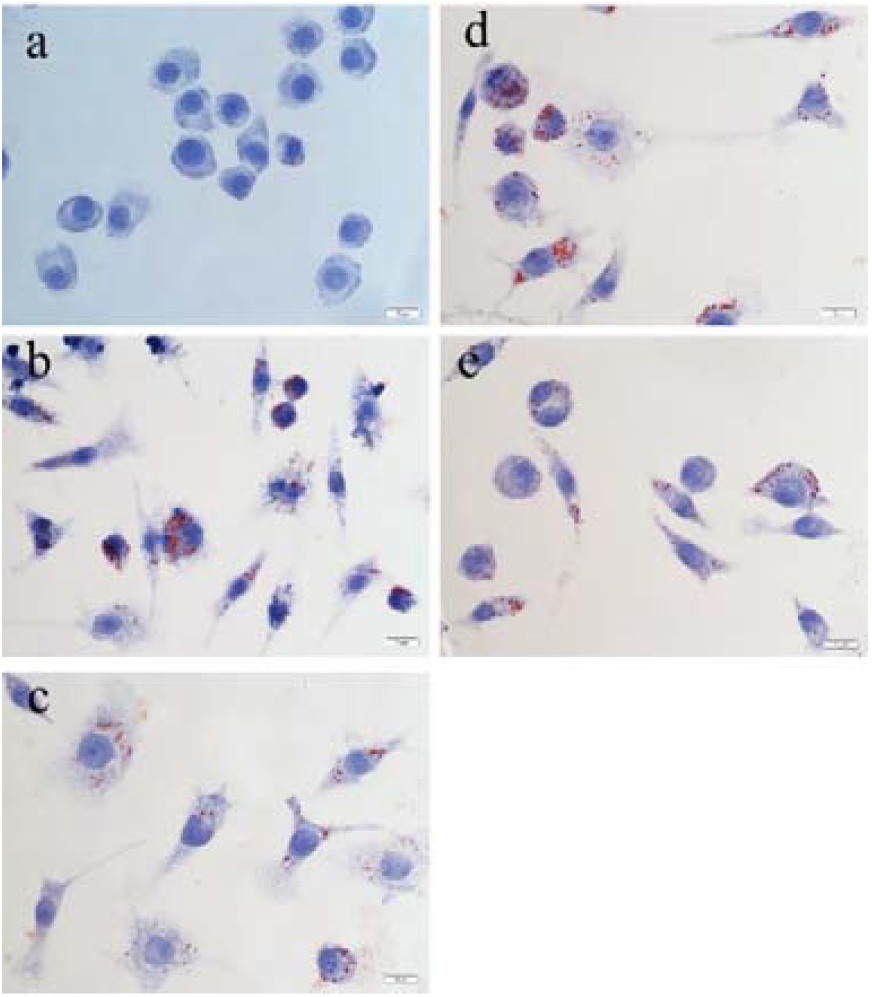
Detection of lipid changes by oil red O **a**: Normal group; **b**: Infection group; **c**: miR-106b-5p inhibitor group; **d**: miR-106b-5p inhibitor+ Creb5-siRNA group; **e**: NC group

**Figure 6J-figure supplement 1.**
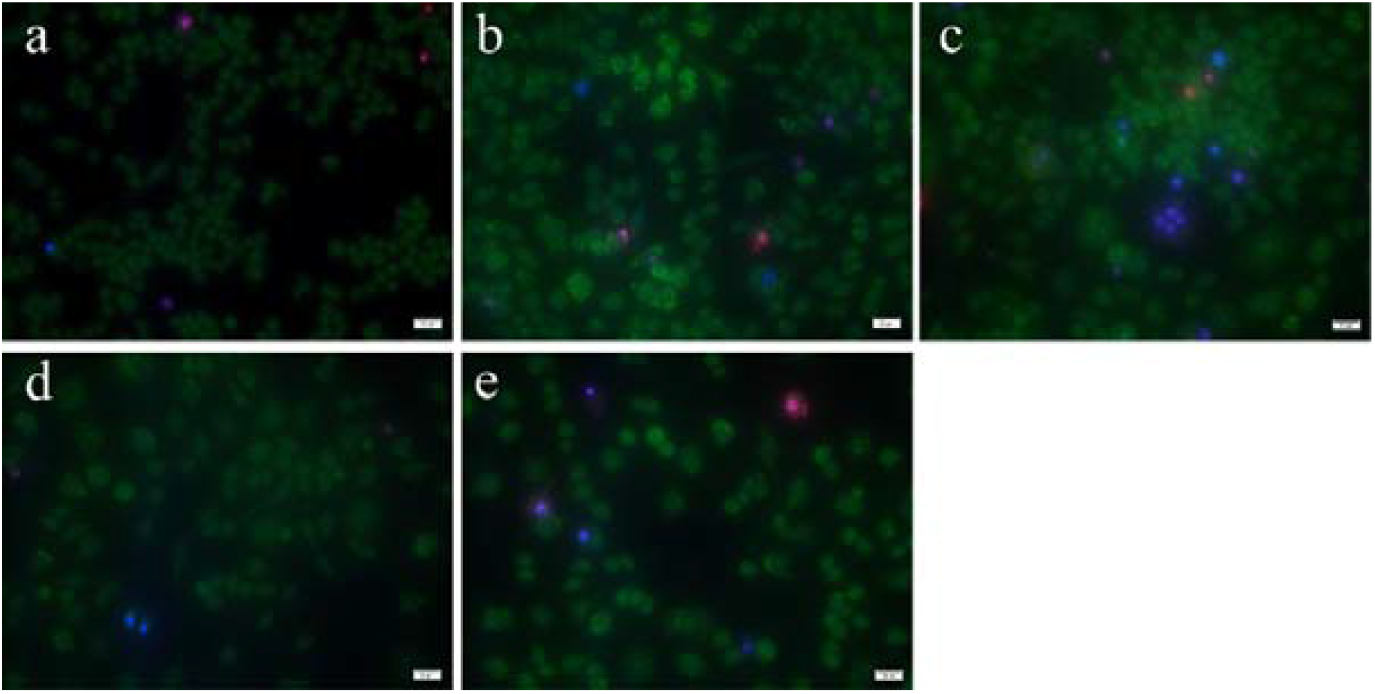
Apoptosis detected by Hoechst/PI **a**: Normal group; **b**: Infection group; **c**: miR-106b-5p inhibitor group; **d**: miR-106b-5p inhibitor+ CREB5-siRNA group; **e**: NC group

